# Uncovering a role for the dorsal hippocampal commissure in episodic memory

**DOI:** 10.1101/415158

**Authors:** M Postans, GD Parker, H Lundell, M Ptito, K Hamandi, WP Gray, JP Aggleton, TB Dyrby, DK Jones, M Winter

**Affiliations:** Cardiff University Brain Research Imaging Centre (CUBRIC), School of Psychology, Cardiff University, Cardiff, UK; School of Psychology, Cardiff University, Cardiff UK; Experimental MRI Centre (EMRIC), School of Biosciences, Cardiff University, Cardiff, UK; Danish Research Centre for Magnetic Resonance, Centre for Functional and Diagnostic Imaging and Research, Copenhagen University Hospital Hvidovre, Hvidovre, Denmark; School of Optometry, University of Montreal, Montreal, Canada and Department of Neurology and Neurosurgery, Montreal Neurological Institute, Montreal, Canada; The Alan Richens Welsh Epilepsy Centre, Department of Neurology, University Hospital of Wales, Cardiff UK; Institute of Psychological Medicine and Clinical Neurosciences, School of Medicine, Cardiff University, Cardiff, UK; Dept of Neurosurgery, Neurosciences Division, University Hospital Wales, Cardiff, UK; Brain Repair And Intracranial Neurotherapeutics (BRAIN) Unit, School of Medicine, Cardiff University, Cardiff, UK; Department of Applied Mathematics and Computer Science, Technical University of Denmark, Kongens, Lyngby, Denmark; School of Psychology, Australian Catholic University, Melbourne, Australia; Department of Clinical Neuropsychology, University Hospital of Wales, Cardiff, UK

**Keywords:** hippocampal commissure, tractography, episodic memory, familiarity, recollection

## Abstract

The dorsal hippocampal commissure (DHC) is a white matter tract that provides inter-hemispheric connections between temporal lobe brain regions. Despite the importance of these regions for learning and memory, there is scant evidence of a role for the DHC in successful memory performance. We used diffusion-weighted MRI (DW-MRI) and white matter tractography to reconstruct the DHC across both humans (in vivo) and nonhuman primates (ex vivo). Across species, our findings demonstrate close consistency between the known anatomy and tract reconstructions of the DHC. Anterograde tract-tracer techniques also highlighted the parahippocampal origins of DHC fibers in nonhuman primates. Finally, we derived Diffusion Tensor MRI (DT-MRI) metrics from the DHC in a large sample of human subjects to investigate whether inter-individual variation in DHC microstructure is predictive of memory performance. The mean diffusivity of the DHC was correlated with performance in a standardised episodic memory task; an effect that was not reproduced in a comparison commissure tract – the anterior commissure. These findings highlight a role for the DHC in episodic memory, and our tract reconstruction approach has the potential to generate further novel insights into the role of this previously understudied white matter tract in both health and disease.

## Introduction

The two hemispheres of the brain are connected by commissural fiber systems that include the corpus callosum, anterior commissure (AC), posterior commissure, ventral hippocampal commissure (VHC), and dorsal hippocampal commissure (DHC) (Demeter et al. 1985). The DHC (alternatively the ‘dorsal psaltarium’) provides inter-hemispheric connections between functionally-related structures in the medial temporal lobes (MTL), including the presubiculum, entorhinal and parahippocampal cortex (Demeter et al. 1985, 1990; Gloor et al. 1993). Given that these regions play a key role in successful learning and memory (Zola-Morgan et al. 1989; Squire and Zola-Morgan 1991; Aggleton and Brown 1999; Aggleton 2012), their ability to communicate effectively with contralateral homologous regions via the DHC may also be important for performance in these cognitive domains.

There have, however, been few studies of the function of the DHC, potentially due to misunderstanding around the cross-species anatomy of the DHC, as distinct from other local fiber populations such as the VHC, fornix, and corpus callosum (Demeter et al. 1985; Raslau et al. 2015; Tubbs et al. 2015). In rodents, the VHC supports dense inter-hemispheric connections between the hippocampi, which originate throughout the long-axis of the hippocampus; in nonhuman primates, VHC connections are reduced so that only the uncal and genual subdivisions of the hippocampal formation are connected to those in the contralateral hemisphere (Demeter et al. 1985; Gloor et al. 1993). By contrast, the DHC remains a substantial tract in nonhuman primates, but it carries commissural projections to and from the parahippocampal region rather than the hippocampus proper. From injection sites in the presubiculum, entorhinal and parahippocampal cortex, tract tracer studies in nonhuman primates have traced labelled DHC fibers into the alveus and fimbria and then the posterior columns of the fornix; these fibers arch dorso-anteriorly and then turn medially to cross the midline along the inferior aspect of the corpus callosum, before taking a mirror-image route back to the contralateral parahippocampal region (Demeter et al. 1985, 1990). Anatomical studies have found no convincing evidence of a VHC in humans but the location of the human DHC corresponds precisely to that reported for nonhuman primates (Gloor et al. 1993). Despite their distinct anatomy, the VHC and DHC are sometimes collectively termed ‘the hippocampal commissure’ (Demeter et al. 1985), and the DHC is sometimes described as part of the fornix (e.g., ‘fornix commissure’) (Mark et al. 1993). It is, however, difficult to infer the function of the DHC from potentially informative clinical case reports and animal studies if it is not appropriately differentiated from these other structures.

In one relevant study highlighting a potential role for the DHC in successful learning and memory, fornix transection did not impair the ability of monkeys to learn concurrent visual discriminations, but the fornix damage in one subject extended to the DHC, and that subject made significantly more errors and required more training sessions to learn the task compared to the slowest control (Moss et al. 1981). This subject was also impaired in an object recognition task (Mahut et al. 1981). Similarly, clinical case reports describe individuals with anterograde amnesia following combined DHC and fornix damage (Heilman and Sypert 1977; D’Esposito et al. 1995), although it is difficult to evaluate the effect of DHC damage in these cases because fornix damage alone is sufficient to produce anterograde amnesia (Aggleton 2008). A deficit in both verbal and visual recall has also been reported in patients who underwent callosotomy surgery for intractable epilepsy but only when the section included the posterior corpus callosum (Clark and Geffen 1989; Phelps et al. 1991). This is pertinent because the rostral splenium and posterior DHC fibers are intermingled, so split brain surgery involving the posterior corpus callosum always involves DHC transection.

The inferences we can derive from these small, methodologically heterogenous studies are, however, limited. Patients with verifiable DHC damage are extremely rare, and there are no reported cases of DHC damage sparing other relevant structures. An alternative approach is to examine whether inter-individual variation in the microstructure of the DHC is related to differences in memory performance. Begré et al., used diffusion tensor imaging (DTI) to search, voxel-wise, for a correlation between a measure of white matter microstructure (inter-voxel coherence) and performance in the Rey Visual Design Learning Test (Begré et al. 2009). In their small sample (N=14), the authors reported that clusters of voxels demonstrating such a relationship overlapped with the DHC. The reported coordinates, however, correspond to the inferior-caudal surface of the splenium, whereas histological studies localise the DHC ventral to the corpus callosum body with posterior DHC fibers becoming intermingled with those of the *rostral* splenium(Demeter et al. 1990). The clusters reported by Begré et al., may therefore lack specificity to the DHC. Wei et al., recently demonstrated that white matter tractography and DW-MRI, can be used to isolate and reconstruct the trajectory of the human DHC, *in vivo*, but no individual subject-level reconstructions were shown (a group-level reconstruction was provided) and the study did not investigate the relationship between DHC microstructure and cognitive performance (Wei et al. 2017). A study in a larger sample is therefore required to isolate the human DHC systematically and investigate the functional role of this tract in memory. Evidence that the DHC can be reconstructed accurately in nonhuman primates, where the tract morphology has been well characterised, would reinforce confidence in the accuracy of human DHC reconstructions.

In the present study, we report a semi-automated tractography approach that can be used to reconstruct the DHC across humans (in vivo) and nonhuman primates (ex vivo). We also present tract tracer findings highlighting that primate DHC fibers form a distinct tract and originate in the parahippocampal region rather than the hippocampal formation. Finally, we derived diffusion tensor imaging metrics from the DHC in a large sample of 100 human subjects, to investigate whether inter-individual variation in the microstructure of this tract correlates with memory performance. We also assessed whether the bilateral volumes of several relevant gray matter MTL regions relates to memory performance.

## Materials and Methods

### Data

#### Ex vivo non-human primate MR data

Diffusion- and T1-weighted MR data that were obtained previously from the perfusion-fixed brains of four healthy adult female vervet monkeys (*Chlorocebus sabeus;* specimens e3429, e3487, e3494, and e4271), were available for analysis (age range = 32-48 months; mean = 36.25, SD = 7.85). The animals were obtained from the Behavioral Science Foundation, St. Kitts and were socially housed in enriched environments. The experimental procedures were reviewed and approved by the Institutional Review Board of the Behavioral Science Foundation, acting under the auspices of the Canadian Council on Animal Care. The postmortem brains were prepared for data collection on a preclinical 4.7 T Agilent scanner system at the Danish Research Centre for Magnetic Resonance using an *ex vivo* imaging protocol reported previously (Dyrby et al. 2011, 2014). This included a DW-MRI pre-scan of at least 15 hours duration to avoid introducing short-term instabilities into the final DW-MRI datasets (e.g., due to motion caused by physical handling of the tissue (Dyrby et al. 2011, 2013)). The brain specimens were also stabilized to room temperature prior to scanning, and a conditioned flow of air around the specimen was maintained throughout scanning to reduce temperature drifts of the diffusion signal (Dyrby et al. 2011, 2013).

Diffusion-weighted images were collected using a diffusion-weighted pulsed gradient spin echo sequence with single line readout. The scan parameters were as follows: repetition time, TR = 7200 (but TR = 8400ms for subject e4271); echo time, TE = 35.9ms; gradient separation, DELTA = 17.0ms; gradient duration, delta = 10.5ms; gradient strength, g = 300 mT/m; number of repetitions, NEX = 2 (averaged offline); matrix size = 128 × 256 with 100 axial slices offering whole brain coverage with isotropic 0.5mm voxels. Gradients were applied along 68 uniformly distributed directions with a b-value of 9686 s/mm^2^ using scheme files available from the Camino tool kit(Cook et al. 2006). Thirteen non diffusion-weighted images with b = 0 s/mm^2^ were also acquired. T1-weighted images were acquired using a 3D MPRAGE sequence with 0.27mm isotropic voxels and the following parameters: TR = 4ms; TE = 2ms; TI = 800ms; FA = 9°; matrix = 256 × 256 × 256, axial image plane.

#### Ex vivo non-human primate anterograde tract tracer data

To highlight the distinct origins of fibers comprising the DHC and the nearby fornix, we examined ex vivo brain specimens obtained from two male cynomolgus monkeys (*Macaca fascicularis* - ACy14 and ACyF23) aged 1-2 years, that had received anterograde tract tracer injections in different medial temporal lobe regions for a previous study of the origin and topography of the fibers comprising the fornix (Saunders and Aggleton 2007). Like the vervet monkeys used for our ex vivo tractography analyses, cynomolgus monkeys are members of the *Cercopithecinae* subfamily of Old World monkeys, and the anatomy of the brain is considered to be very similar across these species (Woods et al. 2011). The reader is referred to the original manuscript for a detailed description of the stereotactic surgery and subsequent brain extraction protocols (Saunders and Aggleton 2007), but briefly, a cocktail of tritiated amino acids was injected into distinct target regions within the medial temporal lobe across the two cases. This cocktail was composed of an equal-parts mixture of either tritiated proline and leucine (final concentration of 50 μCi/μl, New England Nuclear), or tritiated proline, leucine, lysine, and an amino acid mixture derived from algal protein hydrosylate (final concentration of 100 μCi/μl, New England Nuclear), and was injected into the surgically exposed target region using a Hamilton syringe. Specimen ACy14 received an injection in the hippocampal formation, centred in the subiculum, whereas specimen ACyF23 received an injection that incorporated the caudal perirhinal cortex and the rostral parahippocampal cortex. Following a 5-10 day postoperative survival period, the two monkeys were deeply anesthetized and the brain was extracted and cryoprotected. The specimens were cut into 33μm coronal sections, coated with emulsion and subsequently exposed at 4 °C for 6-30 weeks before being developed and counterstained for thionine. Specimen ACyF23 had undergone a bilateral fornix transection procedure 2-12 months prior to the injection of the tritiated amino acids; the case is nevertheless informative because the subsequent amino acid injections resulted in labelling up to the point of the transection.

#### In vivo human MR and cognitive data

Cognitive, diffusion- and T1-weighted MR data were obtained for 100 subjects from the Q3 release of the Human Connectome Project (50 males, aged 22-35 years) (Glasser et al. 2013; Sotiropoulos et al. 2013; Van Essen et al. 2013). The participants in that previous study were recruited from Washington University and the surrounding area and gave informed consent in line with policies approved by the Washington University Institutional Review Board.We co-opted these data for the present analyses to exploit the high-quality diffusion-weighted images that are acquired through the HCP owing to the superior gradient strengths afforded by their customized gradient set. This subsample of the available HCP data will henceforth be referred to as the ‘HCP dataset’.

For each subject, whole-brain diffusion- and T1-weighted images had been acquired on a customized 3T Connectom Skyra scanner (Siemens, Erlangen) with a 32-channel head coil and a customised SC72C gradient set. Each pre-processed dataset comprised 90 diffusion directions for each of three shells with *b*-values of 1000, 2000 and 3000 s/mm^2^; these images were acquired with TR = 5500ms, TE = 89ms and 1.25mm^3^ isotropic voxels. 18 images with *b* = 0 s/mm^2^ were also acquired. Corresponding T1-weighted images were acquired by taking two averages using the 3D MPRAGE sequence (Mugler and Brookeman 1990), with 0.7mm^3^ isotropic voxels and the following parameters: TR = 2400ms, TE = 2.14ms, TI = 1000ms, FA = 8°, FOV = 224mm, matrix = 320 × 320 × 256 sagittal slices in a single slab. Note that the pre-processed HCP diffusion datasets are aligned to the T1 images using FLIRT (Jenkinson and Smith 2001; Jenkinson et al. 2002) as standard so that both the diffusion and T1 data that were available to us were pre-aligned in 1.25mm native structural space. Further acquisition parameters and details of the minimal MR pre-processing pipeline have been reported previously (Glasser et al. 2013; Sotiropoulos et al. 2013).

Available cognitive data for the HCP subjects included performance in the Computerized Penn Word Memory task (CPWM)(Moore et al. 2015), the Picture Sequence Memory Test (PSMT) (Dikmen et al. 2014) and the List Sorting Working Memory Test (LSWMT) (Tulsky et al. 2014). The CPWM is a verbal episodic memory task in which a participant is required to discriminate 20 pre-exposed target word stimuli from 20 inter-mixed novel distractor stimuli; performance is quantified here as subjects’ total number of correct responses. In the PSMT, another episodic memory task, subjects are required to learn and recall a sequence of picture stimuli over a number of trials and performance is scored as the cumulative number of adjacent pairs of pictures that are correctly recalled over 3 learning trials. In the LSWMT, subjects are presented with a series of picture stimuli on a computer screen (e.g., an elephant and a mouse) and are required to remember the stimuli comprising the sequence, mentally reorder them from smallest to largest, and finally recite the revised sequence of stimuli; performance is scored as the number of correct responses across the stimulus lists that comprise this working memory task. For both the PSMT and LSWMT, HCP subjects’ raw scores have been standardised against the NIH Toolbox normative sample (Weintraub et al. 2013). These standardised scores can also be age-adjusted, but given that we had non age-adjusted raw scores for the CPWM, we used subjects’ unadjusted PSMT and LSWMT scores for subsequent analyses.

Finally, pre-existing regional volume measures were available for a number of relevant cortical and subcortical regions in the HCP dataset, because the T1- and T2-weighted images that are acquired for the HCP are segmented using Freesurfer software as part of the standard pre-processing pipeline (Glasser et al. 2013). We used these data to investigate whether differences in CPWM, PSMT and/or LSWMT performance, are also related to the volume of several key gray matter regions within the MTL. Our specific regions-of-interest (ROIs) were the hippocampi, amygdalae, entorhinal cortex, parahippocampal cortex (areas TH and TF), as well as the temporal pole, and estimates of total Intracranial Volume (ICV). The hippocampus was of interest because the DHC is sometimes assumed to support dense inter-hippocampal connections, despite an absence of confirmatory evidence (Demeter et al. 1990). By contrast, the entorhinal and parahippocampal cortices are known to project to contralateral structures via the DHC (Demeter et al. 1985, 1990), and they also provide a functionally important input/output pathway for the hippocampus itself (Aggleton 2012). The temporal pole and amygdala were ROIs that are known to project to or receive from contralateral structures via the anterior commissure (AC), which was used as a comparison tract for our tractography analyses, as described below (Klingler and Gloor 1960; Turner et al. 1979; Demeter et al. 1985).

### Data Processing

#### Ex vivo non-human primate MR data

The T1-weighted images for each nonhuman primate specimen were masked to contain only brain tissue using FSL utilities (Smith et al. 2004). Gray/white matter contrast is reversed in our T1 images of *ex vivo* tissue (Dyrby et al. 2018); we therefore inverted the T1-weighted images for subsequent processing and display purposes. The T1-weighted brain images were then submitted to the standard Nonhuman Primate EMSegmenter pipeline in 3DSlicer version 4.9.0. The pipeline registers the T1-weighted image to a probabilistic vervet monkey MRI atlas using BRAINSFit (Johnson et al. 2007; Fedorov, A., Li, X., Pohl, K.M., Bouix, S., Styner, M., Addicott, M., Wyatt, C., Daunais, J.B., Wells, W.M., & Kikinis 2011), and segments the image into unilateral ROIs, including the hippocampus, using the EM segmenter algorithm (Pohl et al. 2007). The subject-specific aligned and unbiased hippocampus segmentations were thresholded at 40%, binarised, and brought into native diffusion space using FLIRT, ready for use as ROIs for tractography.

Visual inspection of the DW-MRI datasets revealed that no additional pre-processing was required to adjust for motion or eddy currents prior to streamline reconstruction (Dyrby et al. 2014). A multiple-ROI tractography approach (see Fig 1*A*) was used to reconstruct the DHC. Tractography was performed from all voxels in the left hippocampus ROI in subjects’ native diffusion-space in ExploreDTI v4.8.3 (Leemans et al. 2009)using a deterministic tractography algorithm based on constrained spherical deconvolution (Tournier et al. 2008; Jeurissen et al. 2011). The contralateral hippocampal ROI was used as an ‘AND’ gate to capture any propagated streamlines that terminated in the contralateral hippocampal/parahippocampal region. Three additional ‘NOT’ ROIs were manually drawn to exclude streamlines corresponding to other pathways. These included: 1) An ROI covering the entire section, drawn on the most inferior axial slice where the body of the corpus callosum was visible, 2) A coronal ROI covering the entire section placed at a slice where the parahippocampal cingulum begins to descend behind the splenium, and 3) A coronal ROI covering the entire section except the temporal lobes, placed at the slice where the anterior fornix columns descend towards the mammillary bodies. Additional exclusionary ROIs were used to remove extant spurious streamlines as required. A step size of 0.1mm and an angle threshold of 60 degrees was applied to prevent the reconstruction of anatomically implausible streamlines. Tracking was performed with a supersampling factor of 4 × 4 × 4, so that streamlines were initiated from 64 grid points, uniformly distributed within each voxel.

**Figure 1.**
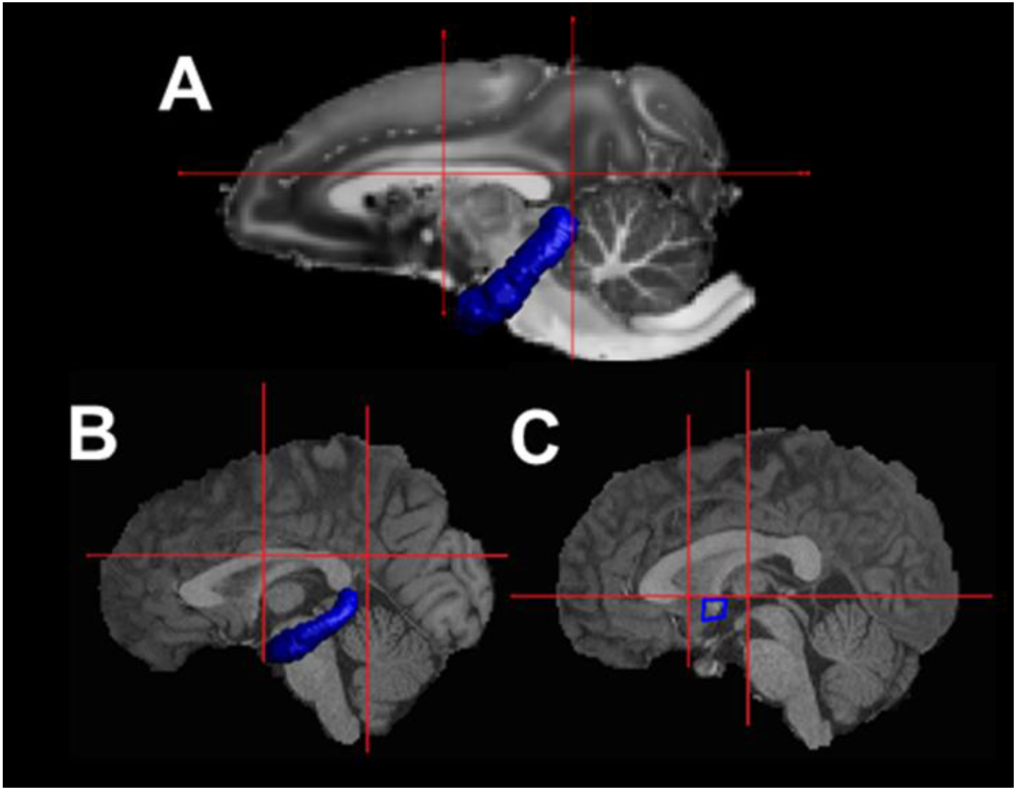
Regions of interest (ROIs) used for dorsal hippocampal commissure (DHC) and anterior commissure (AC) tractography. The hippocampal (blue) and manually drawn (red lines) ROIs used for DHC tractography shown on a mid-sagittal section of a T1-weighted image for a representative ex vivo nonhuman primate specimen in 0.5mm^3^ native diffusion space (A) and a Human Connectome Project (HCP) subject in 1.25mm^3^ native diffusion space (B). Also shown are the manually-drawn ROIs used for AC tractography (red and yellow lines) in a representative HCP subject (C).

#### Ex vivo non-human primate anterograde tract tracer data

A Leica DM500B microscope with a Leica DFC310FX digital camera and Leica Application Suite v4.7 image acquisition software were used to obtain both bright- and dark-field images from our ex vivo cynomolgus monkey specimens.

#### In vivo human MR and cognitive data

Whole brain voxel-wise maps of two DTI measures of white matter microstructure – Fractional Anisotropy and Mean Diffusivity (FA and MD, respectively) (Basser and Pierpaoli 2011) - were derived from the *b* = 1000 s/mm^2^ images. Unilateral hippocampal ROIs were segmented from subjects’ T1-weighted images using FIRST (Patenaude et al. 2011). Streamlines were then seeded from the left hippocampus using the same combination of ROIs described above (Fig 1*B*), and a multi-shell multi-tissue constrained spherical deconvolution algorithm (MSMT-CSD) was applied to the subjects’ complete diffusion dataset (Jeurissen et al. 2014). This process was then repeated with tractography seeded from the right hippocampus. Tracking parameters were the same as above except a step size of 0.5 mm was applied. Two additional ROIs were then drawn around the DHC reconstructions on sagittal sections located 5 slices from the midline of the brain to extract a transverse segment of the DHC, where the streamlines are well differentiated from those of other local white matter pathways. This was done separately for the reconstructions obtained by seeding tractography from the left and right hemispheres. The transverse DHC segments were intersected with the whole brain voxel-wise FA and MD maps. For both FA and MD, the mean measures obtained from the two segments were then combined into a vertex-weighted mean measure as follows:

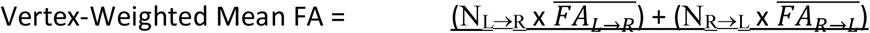

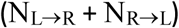

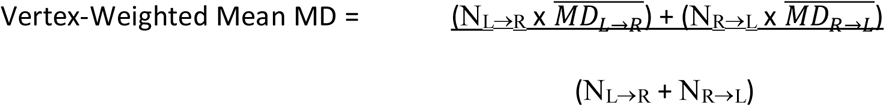

where N_L→R_ and N_R→L_ refer to the number of vertices comprising the tract segment obtained by seeding tractography from the left and right hemisphere, respectively. 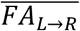 and 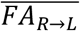 refer to the mean FA measure obtained from the left- and right-seeded segment; likewise, 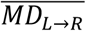 and 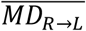 refer to the mean MD measure obtained from the left- and right-seeded segment. These vertex-weighted measures of mean FA and MD take into account any potential differences in the number of streamlines that comprise the left versus right-seeded segments, and were later correlated with memory measures. For the sake of brevity, in the remainder of the text we refer simply to mean FA and MD measures without reference to the vertex-weighting that was applied.

For comparison, these measures were also obtained from a transverse segment of the anterior commissure (AC). The AC is a commissural fiber pathway – the function of which is not well understood – that provides interhemispheric connections between the temporal pole, the amygdala, the superior and inferior temporal gyri, and the parahippocampal gyrus (Demeter et al. 1990). Given that both the DHC and AC contain fibers that originate and cross in the parahippocampal gyrus, we restricted our AC analyses to those relatively ‘anterior projections’ of this fiber bundle, which involve the temporal pole and amygdala. This was achieved by seeding tractography from an ROI manually drawn around the AC on a sagittal section 5 slices from the midline, where the AC is visible at the point it bifurcates the descending fornix columns (see Fig 1*C*). This ‘SEED’ ROI was initially drawn in the left hemisphere, and a corresponding ‘AND’ ROI was placed at the same point in the right hemisphere. An exclusionary ‘NOT’ ROI with whole-brain coverage was then drawn on an axial slice immediately above the AC. Another, covering the whole brain except the temporal lobes, was drawn on a coronal section immediately posterior to the rostrum of the corpus callosum. A final ‘NOT’ ROI was drawn around the whole brain on a coronal section located just anterior to the pons. This procedure was repeated with the seed and the AND ROIs placed in the opposite hemispheres. The initial seed and ‘AND’ ROIs were then used to extract a transverse segment of the AC from both reconstructions. Mean FA and MD metrics were extracted and combined using the above formula.

#### Statistical Analysis

Two-tailed Pearson correlation statistics were used to investigate the relationship between DT-MRI measures of DHC and AC microstructure (FA and MD), and performance in three standardised memory tasks (CPWM, PSMT and LSWMT). Correlation statistics were computed with 1000 bootstrapped samples to derive 95% confidence intervals, and a Bonferonni-Holm step-down procedure was used to adjust derived p-values for six structure-cognition correlations, separately for each tract (the DHC and AC). Subjects in whom both the DHC and AC were successfully reconstructed were included in these analyses to enable fair comparisons between dependent correlations across these tracts. To test for differences between correlations across the DHC and AC, any significant structure-cognition associations identified in one tract, were compared with the corresponding correlation in the other tract using one-tailed Steiger Z tests, which are reported alongside Cohen’s *q* effect size measures (Cohen 1988). A significance threshold of *p* = 0.05 was used for all comparisons.

The same correlational approach was used to investigate the relationship between memory performance (in the CPWM, PSMT and LSWMT) and the volume of individual temporal lobe regions including the amygdala, hippocampus, temporal pole, entorhinal and parahippocampal cortices. Bilateral volume measurements were used to maximise statistical power, and *p*-values were Bonferonni-Holm adjusted for fifteen volume-cognition correlations. Regional gray matter volume measures are often confounded by inter-individual differences in total intracranial volume (ICV); we therefore used the following formula to adjust volume measurements for differences in ICV prior to any correlational analyses:

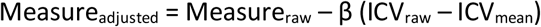

where ICV_raw_ refers to a subject’s ICV estimate, ICV_mean_ refers to the mean ICV in the HCP dataset, and β refers to the slope of the regression line between ICV and the measure of interest (Voevodskaya et al. 2014).

## Results

### Ex vivo non-human primate MR data

To demonstrate the feasibility of a white matter tractography approach for investigating the role of the DHC in human episodic memory, we first applied a multiple region-of-interest (ROI) deterministic tractography protocol to diffusion- and T1-weighted images obtained from four ex vivo nonhuman primate brain specimens (see Methods, Fig 1 and Fig 2*A*). In all four specimens, this revealed a large number of streamlines (mean: 2367.75, SD: 1383.791) that were broadly consistent with the known anatomy of the non-human primate DHC (see Fig 2*B-E*). At the midline of the brain, the transverse portion of the DHC streamlines was situated along the anterior and inferior aspect of the rostral splenium of the corpus callosum. More laterally, these streamlines arched inferiorly towards the hippocampus and parahippocampal region. Whilst a number of these streamlines progressed inferiorly towards regions along the parahippocampal gyrus (see Fig 2*A*, right), consistent with the known anatomy, a number terminated in or around the hippocampus after having intersected our hippocampal ROIs. This finding highlights limitations in resolving crossing fiber populations with existing tractography techniques, and the fact that near the tail of the hippocampus, DHC fibers are known to merge with those of the fornix-fimbria and the alveus, which then cover the hippocampus (Gloor et al. 1993). Indeed, Fig 3*A-C* shows the DHC reconstruction for a representative specimen alongside streamlines corresponding to the fornix, which were reconstructed for illustrative purposes using a multiple-ROI approach reported previously (Metzler-Baddeley et al. 2011); whilst the transverse portion of the DHC is readily differentiated, more laterally, many of the DHC streamlines become intermingled with those of the fornix as the latter covers the hippocampus. Nevertheless, these ex vivo DHC reconstructions suggest that white matter tractography can be used to detect and reconstruct inter-hemispheric DHC connections, and that the transverse portion of these fiber pathway reconstructions in particular, is well characterised and differentiated from the fornix. The reconstructions were similar in humans (Fig 3*D*), so our subsequent quantitative analyses in human subjects were based on mean microstructure measures that were extracted from this transverse portion of the DHC (see Methods).

**Figure 2.**
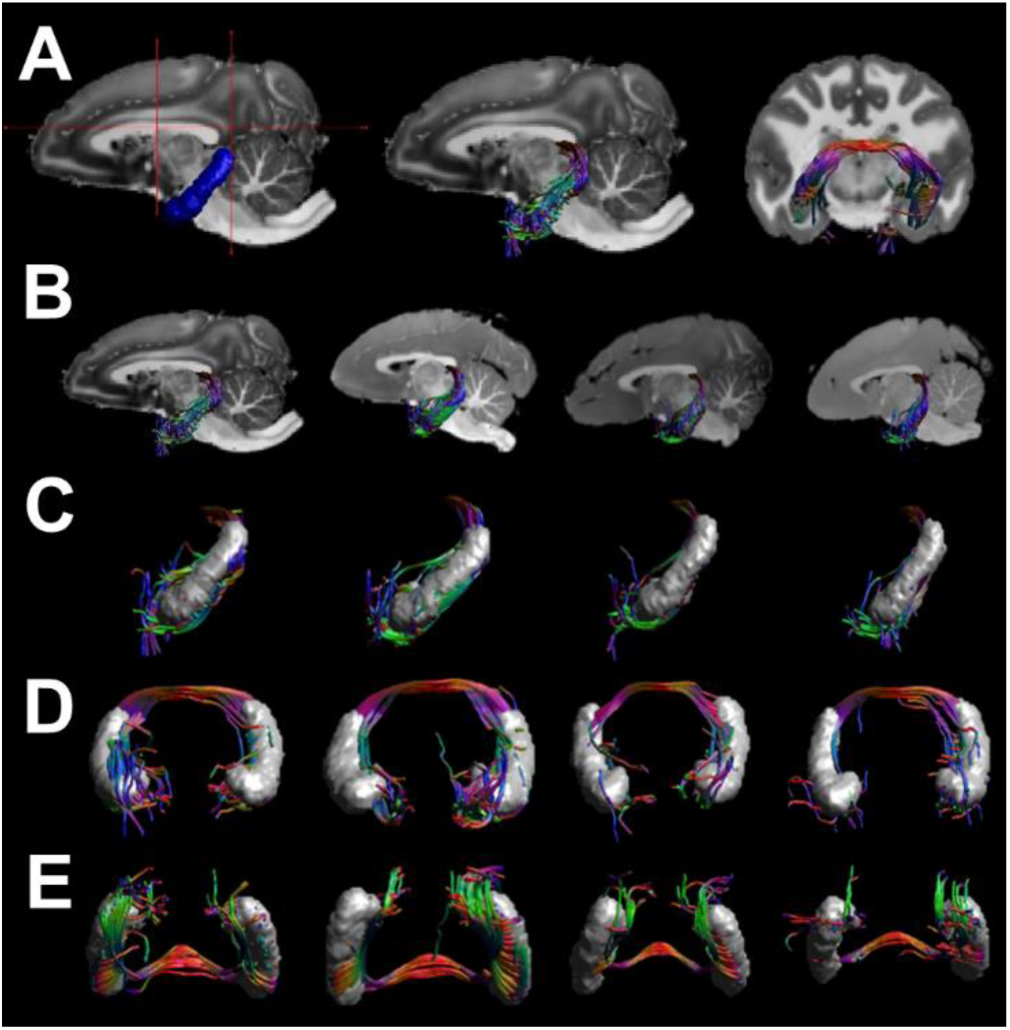
Regions-of-interest (ROIs) used to extract the dorsal hippocampal commissure (DHC) and the subsequent tract reconstructions. The hippocampal (blue) and manually-drawn (red lines) ROIs used to extract the DHC in one representative specimen, and the subsequent reconstructions shown over a mid-sagittal and coronal section from the corresponding T1-weighted image in 0.5mm^3^ native diffusion space (A); The reconstructions in all four specimens (B); The DHC reconstructions illustrated from a left-lateral, anterior-posterior, and inferior-superior perspective, alongside the anatomical hippocampal ROIs for spatial context (C, D and E, respectively). Note that for computational purposes, these renderings contain a 1/8^th^ subsample of all reconstructed streamlines.

**Figure 3.**
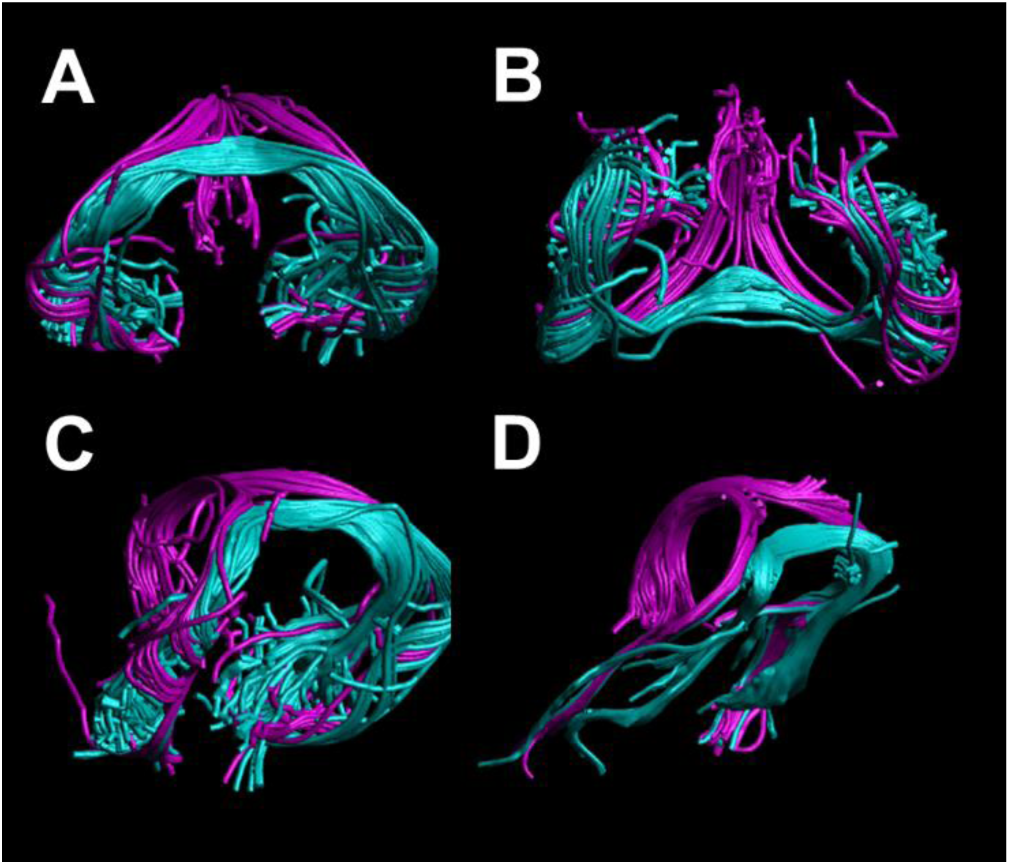
Dorsal hippocampal commissure (teal) and fornix (purple) streamlines reconstructed in representative cases. Streamlines corresponding to these tracts are shown for a representative ex vivo nonhuman primate specimen, shown from a rear coronal-oblique (A), ventral (B) and left-lateral oblique (C) perspective. For comparison, streamlines corresponding to these two tracts are also shown for a representative Human Connectome Project (HCP) subject, from a left-lateral oblique perspective (D).

### Ex vivo non-human primate tract tracer data

To highlight the veridical cortical origins of the inter-hemispheric DHC streamlines that were reconstructed in the previous analysis, we next examined bright- and dark-field photomicrographs taken from two ex vivo nonhuman primate specimens that had previously received anterograde tract tracer injections of radioactive amino acids in different locations within the medial temporal lobe. In a specimen that received an injection in the hippocampal formation itself, centred on the subiculum, strong labelling was present in the ipsilateral but not the contralateral fornix or the DHC (Figs 4*A-C*). This is consistent with previous research showing that the majority of fibers comprising the fornix originate in the subicular cortices and CA subregions of the hippocampal formation, and that neither the fornix or DHC supports inter-hemispheric connections between these regions (Saunders and Aggleton 2007). By contrast, dark field photomicrographs from a specimen that received an injection of tracer into caudal perirhinal and rostral parahippocampal cortex, revealed labelling in both the left and right DHC but almost no label in either the ipsilateral or contralateral fornix (Fig 4*D-E*). This distribution is consistent with DHC fibers originating in regions within the parahippocampal gyrus rather than the hippocampus proper. These findings highlight the distinct cortical origins of the nonhuman primate fornix and DHC.

**Figure 4.**
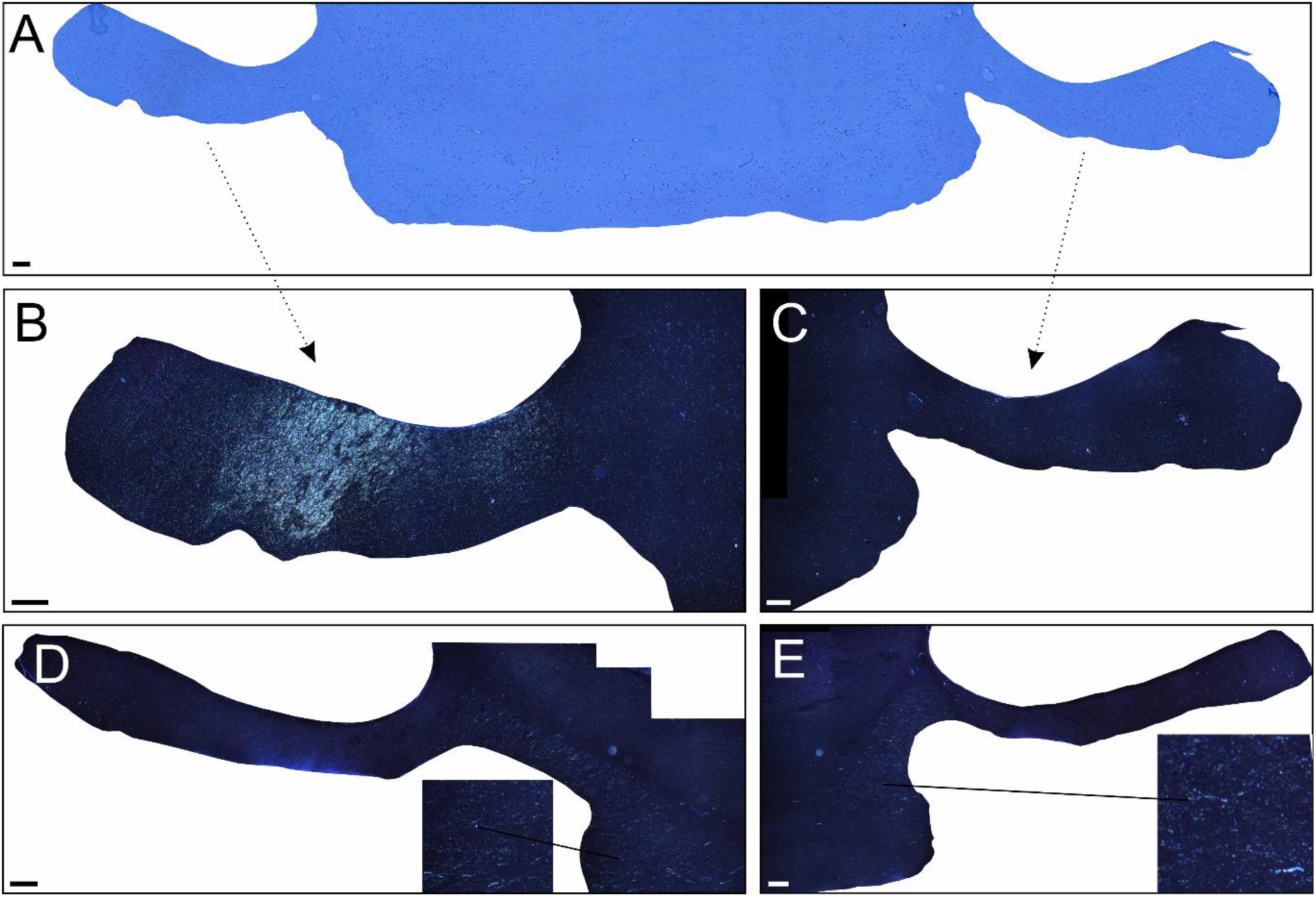
Bright- and dark-field photomicrographs of coronal sections taken at the level of the posterior fornix and dorsal hippocampal commissure (DHC), inferior to the corpus callosum. A) is a bright field photomicrograph from a case which received an anterograde tracer injection in the hippocampal formation centred in the subiculum. B) and C) show dark field photomicrographs of the same section showing strong labelling in the ipsilateral (B), but not contralateral (C), fornix; there was no obvious aggregation of label in the DHC. D) and E) show dark field photomicrographs of a separate section from another case whose injection incorporated caudal perirhinal and anterior parahippocampal cortex; these contain almost no label in the fornix but label in both the left and right DHC. Magnified inserts are included in panels D-E to aid visibility of subtle DHC labelling in the medial portion of the images. Scale bars = 200μm.

### In vivo human MR and cognitive data

#### Association between DHC microstructure and episodic memory performance

We derived diffusion tensor imaging metrics (Fractional Anisotropy, FA, and Mean Diffusivity, MD (Basser and Pierpaoli 2011) from the DHC in a sub-sample of 100 participants in the Human Connectome Project (HCP) for whom T1- and diffusion-weighted MRI data was available for analysis, along with performance in three standardised memory tasks (the Picture Sequence Memory Test, PSMT (Dikmen et al. 2014); List Sorting Working Memory Test, LSWMT (Tulsky et al. 2014); and Computerized Penn Word Memory task, CPWM (Moore et al. 2015)). This enabled us to investigate whether inter-individual variation in DHC microstructure was correlated with memory performance. For comparison, these analyses were repeated in another commissure tract – the AC (see Methods).

Figure 5 illustrates the DHC reconstructions in four representative HCP subjects. Streamlines broadly consistent with the known anatomy of the DHC were successfully reconstructed in 96 subjects (96%). Similarly, streamlines consistent with AC anatomy were successfully reconstructed in 99 subjects (99%). Measures of DHC and AC microstructure (FA and MD) are reported in Table 1.

**Figure 5.**
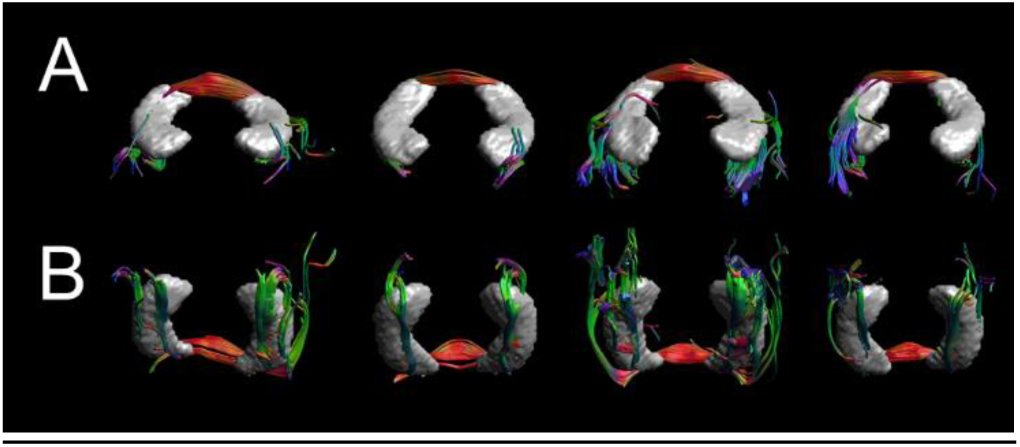
The dorsal hippocampal commissure reconstructions in four representative Human Connectome Project datasets. The reconstructions are shown in the coronal (A) and axial (B) plane.

**Table 1.** Mean measures of DHC and AC microstructure in the HCP dataset (FA and MD) and N streamlines reconstructed. Standard deviations are provided in brackets.

| Tract | Mean FA | Mean MD ($\times 10^{-3} \text{mm}^2 \text{s}^{-1}$) | Mean N Streamlines |
| --- | --- | --- | --- |
| DHC | 0.318 (0.059) | 1.478 (0.163) | 1622.47 (1981.16) |
| AC | 0.439 (0.05) | 0.854 (0.051) | 4102.68 (1896.59) |

Cognitive performance in the CPWM, PSMT and LSWMT is reported in Table 2. A series of two-tailed Pearson correlation analyses revealed a significant negative association between MD and CPWM performance in the DHC (*r* = -0.269, *p* = 0.048, 95% CI = [-0.499, -0.017]), which was not evident in the AC (*r* = 0.100, *p* = 1.0, 95% CI = [-0.123, 0.297]); further, these correlations were significantly different from one another (*Z* = -2.608, *p* = 0.009, *q* = 0.376; see Fig 6). The correlations between DHC MD and both PSMT and LSWMT performance were not statistically significant (*r* = -0.072, *p* = 1.0, 95% CI = [-0.260, 0.096]; *r* = -0.047, *p* = 0.649, 95% CI = [-0.260, 0.159], respectively); although they were not significantly different from the association between DHC MD and CPWM performance (*Z* = -1.517, *p* = 0.065, *q* = 0.204; *Z* = -1.6, *p* = 0.055, *q* = 0.229, respectively). Across the DHC and AC, there were no other statistically significant structure-cognition associations (largest *r* = 0.211, *p* = 0.240, 95% CI = [0.0, 0.403]). These findings imply a potential role for the DHC in CPWM performance – a standardised episodic memory test.

**Table 2.**
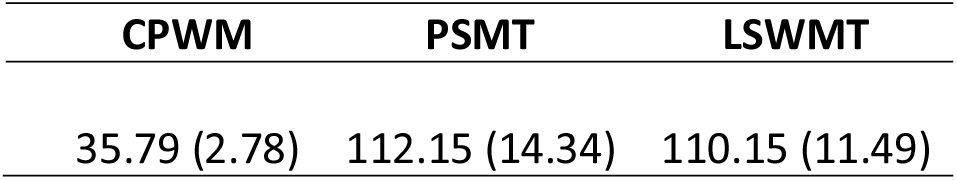
Mean performance in the CPWM, PSMT and LSWMT. Raw scores are reported for the CPWM, and scaled scores are reported for the PSMT and LSWMT. Standard deviations are provided in brackets.

| CPWM | PSMT | LSWMT |
| --- | --- | --- |
| 35.79 (2.78) | 112.15 (14.34) | 110.15 (11.49) |

**Figure 6.**
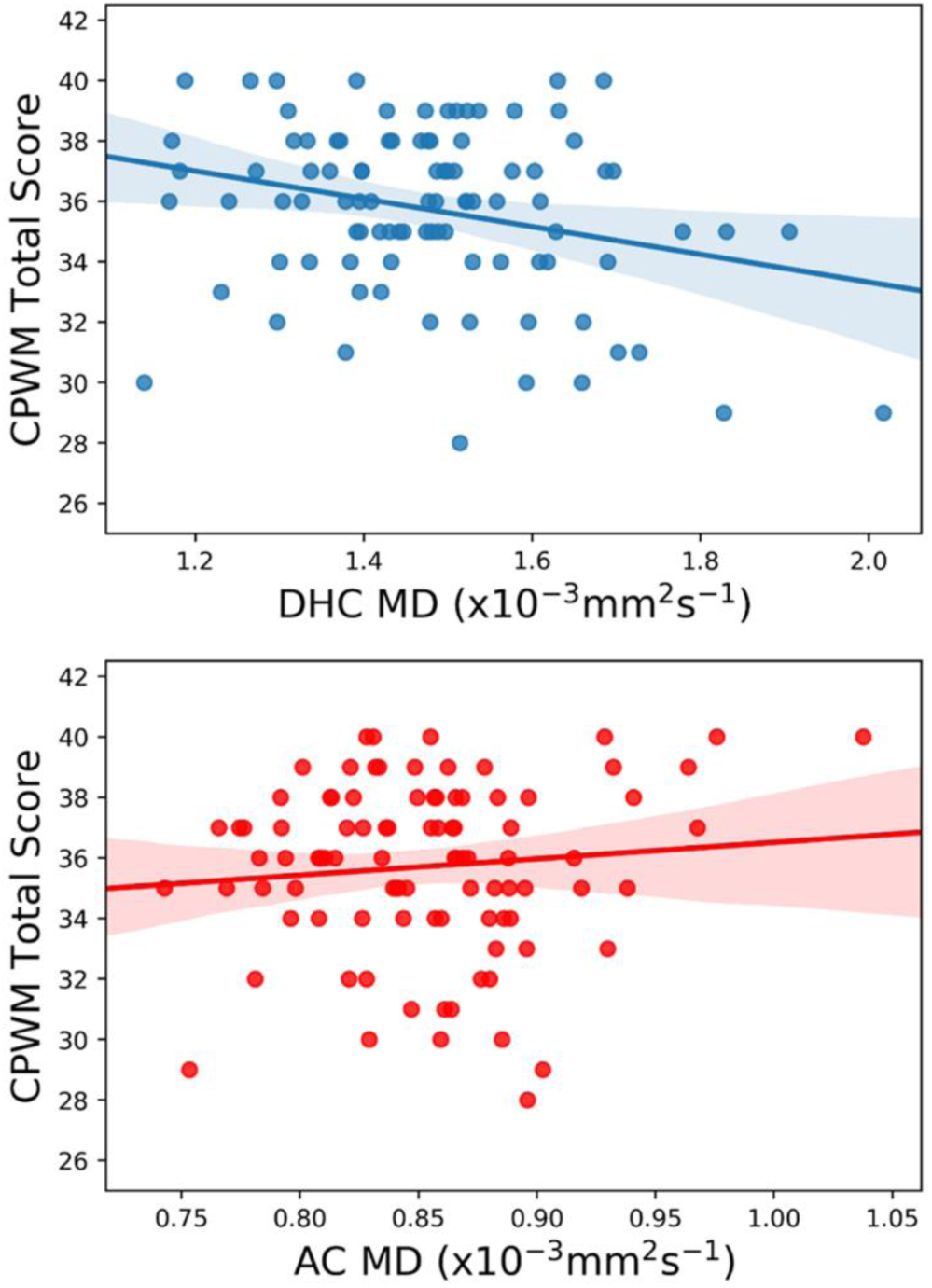
Structure-cognition correlations reported in the text. The correlations between white matter mean diffusivity (MD) and computerized Penn word memory (CPWM) total scores in the dorsal hippocampal commissure (DHC; top) and anterior commissure (AC; bottom). The best fitting linear regression line is plotted alongside 95% confidence intervals.

#### Association between temporal regional volumes and memory

Using pre-existing regional volume estimates for our HCP sub-sample, we assessed whether the bilateral volumes of MTL gray matter regions including the hippocampus, amygdala, temporal pole, entorhinal and parahippocampal cortex, are also related to memory performance. These measures were first adjusted for differences in total Intra-Cranial Volume (ICV; see Methods), and are reported in Table 3. There was no significant association between performance in any of the cognitive tasks (CPWM, PSMT, or LSWMT) and the bilateral ICV-adjusted volumes of these temporal regions (largest *r* = -0.189, *p* = 0.885, 95% CI = [-0.368, -0.004]).

**Table 3.** Mean ICV and ICV-adjusted volumes of bilateral temporal regions. Standard deviations are provided in brackets.

| Structure | Volume (mm<sup>3</sup>) |
| --- | --- |
| Total ICV | 1587548.46 (176651.55) |
| Hippocampus | 8856.07 (663.78) |
| Amygdala | 3210.26 (310.89) |
| Entorhinal Cortex | 3460.21 (575.65) |
| Parahippocampal Cortex | 4391.78 (523.85) |
| Temporal Pole | 4682.52 (490.93) |

## Discussion

This study demonstrated that white matter tractography can be used to reconstruct the DHC in both nonhuman primates (ex vivo) and humans (in vivo), and that these reconstructions broadly conform to the known anatomy of this understudied commissural fiber bundle. That these connections are distinct from those comprising the adjoining fornix is supported by the differential pattern of labelling observed in two ex vivo non-human primate specimens injected with anterograde radioactive tracer in either the subiculum of the hippocampal formation (dense labelling in the ipsilateral fornix) or perirhinal/parahippocampal cortex (labelling in the DHC). Inter-individual variation in the MD of the DHC reconstructions was also correlated with performance in a standardised episodic memory task – the CPWM. Importantly, this structure-cognition association was not evident in another commissural fiber bundle – the AC – implying a degree of specificity in the association between DHC microstructure and CPWM performance. The bilateral volumes of several temporal gray matter regions were not correlated with memory performance.

That our tractography approach affords reconstructions of the DHC across species is consistent with a preservation of DHC morphology across humans and nonhuman primates (Gloor et al. 1993). Wei et al., recently showed that a combination of manually and anatomically-defined ROIs, including the hippocampus, could be used to reconstruct the human DHC in vivo (Wei et al. 2017). Our findings augment those of Wei et al., by demonstrating that this approach is reliable for large datasets, and by clearly defining a specific combination of ROIs that yield DHC reconstructions that broadly reflect the known anatomy. The transverse portion of the DHC reconstructions, in particular, is well differentiated from other local fiber populations such as the fornix, and was therefore the focus of our subsequent quantitative analyses.

The DHC is frequently described as a component of the fornix that supports inter-hemispheric connections between the hippocampi (Raslau et al. 2015; Tubbs et al. 2015). Whilst consistent with the DHC reconstructions shown here, this anatomical interpretation is potentially misleading. Our anterograde tract tracer data, for instance, highlights the distinct hippocampal and parahippocampal origins of the fibers comprising the fornix and DHC in nonhuman primates. The present specimens were previously reported alongside others with more varied injections within the hippocampal formation; again, only in cases where injections extended to parahippocampal regions was any incidental DHC labelling noted (Saunders and Aggleton 2007). The cortical origins of human DHC fibers have not been directly confirmed but the human DHC may also connect parahippocampal regions rather than the hippocampi. Hippocampal ROIs can nevertheless be used to seed DHC tractography, as evidenced by our tract reconstructions, because the hippocampus is covered by the alveus and fimbria-fornix, which do themselves briefly merge with the DHC at the tail of the hippocampus (Gloor et al. 1993). The successful propagation of DHC streamlines from hippocampal ROIs may therefore reflect limitations in resolving crossing fiber populations that are associated with current tractography techniques (Jones et al. 2013). The same limitations result in a proportion of those streamlines also terminating in the contralateral hippocampal ROIs, but several streamlines did nevertheless terminate in contralateral parahippocampal regions. Although the DHC and fornix are partially contiguous, their distinct cortical origins suggests that the former may play a unique – if complementary – role in mnemonic processing.

The association between DHC MD and CPWM performance highlights a role for the DHC in episodic memory. We were, however, limited to analysing cognitive data from the HCP cognitive task battery, which is not necessarily optimised to investigate the role of the DHC in different memory processes. Further interpretation of the relationship between DHC MD and CPWM performance is therefore not straightforward. The CPWM is an episodic memory task in which participants must discriminate between novel and pre-exposed words, but participants are not required to perform free-recall of studied items. According to dual-process models of recognition memory (Aggleton and Brown 1999; Diana et al. 2007; Brown et al. 2010), performance in such tasks could be supported by a *familiarity-based* recognition memory process, which is dependent on parahippocampal regions within the MTL (particularly perirhinal cortex), rather than the hippocampus, which is instead critical for successful performance in tasks that require conscious *recollection* (e.g., free recall). Our findings therefore tentatively suggest that the DHC, may play a role in successful familiarity-based recognition memory.

CPWM performance is not, however, a process-pure measure of familiarity-based recognition memory. Although the CPWM places no explicit demands on recollection, this putatively distinct mnemonic process may also be recruited to aid CPWM performance. Furthermore, the association between DHC MD and CPWM performance was not significantly different to that between DHC MD and PSMT performance, and successful performance in the latter task may be more dependent upon recollection processes. To disentangle the specific memory processes that are partly dependent on DHC connections, future studies should employ a variety of episodic memory paradigms that place differential demands on familiarity and recollection-based recognition memory, including both free-recall and forced-choice recognition tasks.

Whilst the CPWM employs verbal stimuli, the PSMT employs visual stimuli, albeit with additional verbal descriptors. The present association between DHC MD and CPWM performance was not significantly different to that between DHC MD and PSMT performance. Our study did not, therefore, reveal a differential role for the DHC in verbal compared to visual memory. Future research should employ matched verbal and visual memory paradigms to ascertain the extent to which inter-hemispheric mnemonic processing of such stimuli depends upon DHC connections. A considerable body of literature indicates a degree of hemispheric specialisation in visual and verbal processing (Gross 1972; Papanicolaou et al. 2002; Nagel et al. 2013). Whilst we identified an association between DHC microstructure and performance in one verbal episodic memory task, it is possible that DHC connections are particularly important for the mnemonic processing of task-relevant conjunctions of visual and verbal information.

There were no significant correlations between measures of either DHC or AC microstructure and performance in the LSWMT, implying that these tracts are not involved in working memory. However, the correlations between DHC MD and a) CPWM performance and b) LSWMT performance, were not statistically different. Future research should include tasks outside the episodic memory domain – ensuring good variability in outcome measures – to assess any contribution of the DHC to other forms of learning and memory. Our analyses also revealed no associations between memory measures and the bilateral volumes of temporal regions that are known to be connected via the DHC/AC, or play a role in successful recognition memory (Zola-Morgan et al. 1989; Squire and Zola-Morgan 1991; Aggleton and Brown 1999; Aggleton 2012; Ranganath and Ritchey 2012). Further investigations are required to understand the complex relationship between episodic memory performance, white matter microstructure and gray matter macrostructure in this region.

Our results imply that the human DHC is not vestigial, which has implications for the treatment of several neurological conditions. The DHC could, for instance, be incorporated into models of the cognitive impact of resective medial temporal lobe epilepsy surgeries (Trenerry et al. 1993; Dupont 2015). Intracranial EEG studies indicate that a subset of seizures with a medial temporal onset have a pattern of contralateral spread to the hippocampus prior to involvement of contralateral neocortex, potentially via the DHC (Gloor et al. 1993; Rosenzweig et al. 2011). Indeed, whether due to bilateral hippocampal pathology or seizure spread, voxel-based morphometry analyses have identified a cluster of voxels that incorporates the DHC, in which white matter volume is reduced in temporal lobe epilepsy cases with bilateral hippocampal sclerosis compared to healthy controls (Miró et al. 2015). Our tractography protocols offer a complimentary approach to investigating whether DHC microstructure is also compromised in epilepsy cases with mesial temporal sclerosis.

An advantage of our hypothesis-driven tractography approach, is that by constraining our analyses to two commissural tracts, we reduce the risk of reporting both false-positive effects in regions for which we have no specific predictions, and false-negative effects when true structure–cognition relationships are obscured following corrections for large numbers of statistical comparisons. Another advantage of our tractography approach, in which we extract DT-MRI-based microstructural indices that are averaged over a given tract-of-interest, is that it may be more sensitive to subtle microstructural differences that are distributed along the length of the tracts compared with voxel-based methods in which such differences must be clustered in order to detect a significant effect in group-level analyses (e.g., Tract-Based Spatial Statistics) (Smith et al. 2006). Voxel-based methods and metrics that take into account dispersed structural differences could, however, provide complimentary evidence of a role for the DHC in memory. Anatomical Connectivity Mapping (ACM), has recently been proposed as a method of quantifying the strength of connectivity of individual voxels with the rest of the brain (Bozzali et al. 2011). Within a tract, the ACM metric at a given voxel may be sensitive to structural differences further along that tract. Similar to our approach, an average or median ACM measure can also be derived for a given tract-of-interest, and used in subsequent structure-behaviour or structure-cognition correlations (Lyksborg et al. 2014). ACM could potentially provide complimentary evidence of a role for the DHC in episodic memory.

Although DHC MD correlated with CPWM performance, DHC FA was not associated with performance. FA and MD are both affected by multiple axonal properties, including myelination, density, diameter, and configuration as well as partial volume interactions with tract size (Vos et al. 2011; Jones et al. 2013). It is therefore not possible to attribute differences between our FA/MD findings to a single white-matter subcomponent.

In summary, to our knowledge this is the first study to use cross-species anatomical evidence to highlight the DHC as a discrete tract in primates and to systematically reconstruct it using advanced tractography techniques. Reconstructions of the human and nonhuman primate DHC broadly conform to the known anatomy of this tract, affording investigations of the role of the DHC in learning and memory. Indeed, we are also the first to demonstrate a correlation between inter-individual variation in the microstructure of *in vivo* DHC tract reconstructions and differences in a measure of episodic memory performance. Our understanding of the unique role of the DHC in human learning and memory, in both health and disease, is sparse, but the approach described here should advance our knowledge of those aspects of human memory that are partly dependent upon inter-hemispheric processing via the DHC.

## Acknowledgements

DKJ and GDP are supported by a Wellcome Trust Investigator Award (096646/Z/11/Z) and a Wellcome Trust Strategic Award (104943/Z/14/Z). MP* is currently supported by the Medical Research Council (MR/N01233X/1). Data used in the preparation of this work were obtained from the MGH-USC Human Connectome Project (HCP) database (https://ida.loni.usc.edu/login.jsp). The HCP project (Principal Investigators: Bruce Rosen, M.D., Ph.D., Martinos Center at Massachusetts General Hospital; Arthur W. Toga, Ph.D., University of California, Los Angeles, Van J. Weeden, MD, Martinos Center at Massachusetts General Hospital) is supported by the National Institute of Dental and Craniofacial Research (NIDCR), the National Institute of Mental Health (NIMH) and the National Institute of Neurological Disorders and Stroke (NINDS). Collectively, the HCP is the result of efforts of co-investigators from the University of California, Los Angeles, Martinos Center for Biomedical Imaging at Massachusetts General Hospital (MGH), Washington University, and the University of Minnesota.

## Author Contributions

MW developed the initial concept for this study, and the subsequent study design was developed by MP*, DKJ, and MW. Ex vivo vervet monkey MR images were previously acquired by HL, MP, and TBD, who contributed these existing datasets for the present tractography analyses. JA obtained and contributed both bright and dark field images from existing ex vivo cynomolgus monkey specimens. MP*, and MW performed tractography analyses with input from GDP, and under the supervision of DKJ. All statistical analyses were performed by MP* and MW with input from DKJ. All authors provided critical revision of the manuscript, thereby providing important intellectual content.

